# Distinct cryo-EM Structure of α-synuclein Filaments derived by Tau

**DOI:** 10.1101/2020.12.31.424989

**Authors:** Alimohammad Hojjatian, Anvesh K. R. Dasari, Urmi Sengupta, Dianne Taylor, Nadia Daneshparvar, Fatemeh Abbasi Yeganeh, Lucas Dillard, Brian Michael, Robert G. Griffin, Mario Borgnia, Rakez Kayed, Kenneth A. Taylor, Kwang Hun Lim

## Abstract

Recent structural studies of ex vivo amyloid filaments extracted from human patients demonstrated that the ex vivo filaments associated with different disease phenotypes adopt diverse molecular conformations distinct from those in vitro amyloid filaments. A very recent cryo-EM structural study also revealed that ex vivo α-synuclein filaments extracted from multiple system atrophy (MSA) patients adopt quite distinct molecular structures from those of in vitro α-synuclein filaments, suggesting the presence of co-factors for α-synuclein aggregation in vivo. Here, we report structural characterizations of α-synuclein filaments derived by a potential co-factor, tau, using cryo-EM and solid-state NMR. Our cryo-EM structure of the tau-promoted α-synuclein filament at 4.0 Å resolution is somewhat similar to one of the polymorphs of in vitro α-synuclein filaments. However, the N- and C-terminal regions of the tau-promoted α-synuclein filament have different molecular conformations. Our structural studies highlight the conformational plasticity of α-synuclein filaments, requiring additional structural investigation of not only more ex vivo α-synuclein filaments, but also in vitro α-synuclein filaments formed in the presence of diverse co-factors to better understand molecular basis of diverse molecular conformations of α-synuclein filaments.

## Introduction

Aggregation of α-synuclein into amyloid filaments is associated with numerous neurodegenerative diseases including Parkinson’s disease (PD), dementia with Lewy bodies (DLB), and multiple system atrophy (MSA) collectively termed synucleiopathy.^1^ Increasing evidence suggests that the protein aggregates play a key role in the initiation and spreading of pathology in the neurodegenerative diseases.^2–6^ It was shown that α-synuclein aggregates are capable of spreading through the brain and acting as seeds to promote misfolding and aggregation like prion.^7–10^ Although precise molecular mechanisms underlying the neurodegenerative disorders have remained elusive, misfolded α-synuclein aggregates including oligomeric and sonicated fibrillar species exhibit cytotoxic activities.^11^ In addition, injection of preformed filamentous α-synuclein aggregates into mice induced PD-like pathology.^7, 12^ Structural elucidation of filamentous α-synuclein aggregates is, therefore, essential to understanding molecular basis of neurotoxic properties of α-synuclein aggregates and developing therapeutic strategies.

α-synuclein is a 140-residue protein expressed predominantly in the dopaminergic neurons.^13^ The intrinsically disordered protein adopts heterogeneous ensembles of conformations. The diverse conformers in the conformational ensemble might be induced to form distinct amyloid aggregates with different molecular conformations depending on experimental conditions (Figure 1).^10, 14, 15^ Indeed, recent high-resolution structural studies using solid-state NMR and cryo-EM revealed that α-synuclein filaments can adopt diverse molecular conformations under various in vitro experimental conditions.^16–21^ Structural analyses of α-synuclein aggregates seeded by brain extracts from PD and MSA patients suggested that the brain-derived aggregates are heterogenous mixtures of filaments that are distinct from in vitro α-synuclein filaments.^22^ Very recently, high-resolution cryo-EM structures of α-synuclein filaments extracted from MSA and DLB patients were reported.^23^ Interestingly, two types of α-synuclein filaments consisting of two twisting asymmetric protofilaments were observed in MSA filaments extracted from 5 patients. On the other hand, ex vivo DLB filaments were untwisted and morphologically different from those of ex vivo MSA. The structural studies revealed that ex vivo α-synuclein filaments are structurally diverse and quite distinct from those of in vitro α-synuclein filaments produced in buffer, suggesting that diverse co-factors may exist in vivo and induce formation of different α-synuclein filaments.

**Figure 1.**
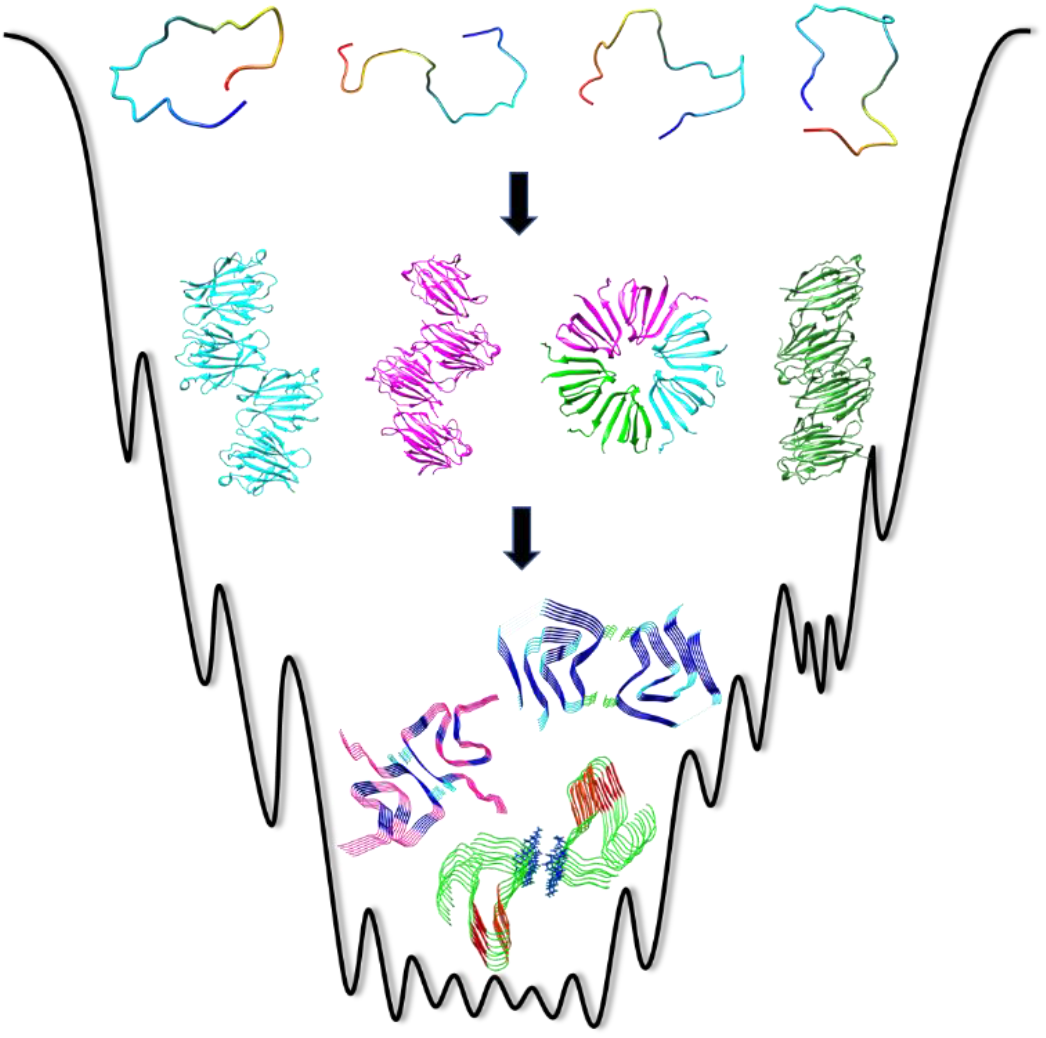
A schematic diagram of energy landscape for α-synuclein aggregation.

α-synuclein remains largely unfolded at low protein concentrations (< 0.1 mM) under physiological conditions. The formation of filamentous aggregates is triggered at aggregation-prone conditions such as higher protein concentrations and more acidic pH.^14, 15^ Misfolding and aggregation of α-synuclein is also promoted by interactions with a variety of co-factors such as lipids, poly(ADP-ribose) (PAR), and other pathological aggregation-prone proteins such as tau and Aβ(1–42) peptides.^14, 24–27^ The co-factors may interact with monomeric α-synuclein and lead to distinct misfolding pathways, resulting in different molecular conformations. Comparative structural analyses of in vitro α-synuclein filaments derived by co-factors and brain-derived ex vivo α-synuclein filaments are required to identify co-factors that promote α-synuclein aggregation in vivo.

Our previous NMR study revealed that tau interacts with the C-terminal region of α-synuclein, accelerating the formation of α-synuclein filaments.^28^ Here we report structural investigation of tau-promoted α-synuclein filaments using solid-state NMR and cryo-EM to investigate the effect of the interactions on the structure of α-synuclein filaments. Our initial solid-state NMR studies indicate that the tau-promoted α-synuclein filaments have similar structural features to those of one of the polymorphs. However, cryo-EM structure of the tau-promoted α-synuclein filaments at 4.0 Å resolution revealed distinct molecular conformations in the N- and C-terminal regions with a much faster helical twist, suggesting that the co-factor, tau, directs α-synuclein into a distinct misfolding and aggregation pathway.

## Experimental Methods

### Protein expression and purification

#### α-Synuclein

Full-length α-synuclein was expressed in BL21(DE3) *E. coli* cells using pET21a plasmid (a gift from Michael J Fox Foundation, Addgene plasmid # 51486) and was purified at 4 ℃ as previously described.^29^ Briefly, the transformed *E. coli* cells were grown at 37 °C in LB medium to an OD_600_ of 0.8. The protein expression was induced by addition of IPTG to a final concentration of 1 mM and the cells were harvested by centrifugation after 12 hrs of incubation at 25 °C. The bacterial pellet was resuspended in lysis buffer (20 mM Tris, 150 mM NaCl, pH 8.0) and sonicated at 4 °C. The soluble fraction of the lysate was precipitated with ammonium sulfate (50%). The resulting protein pellet collected by centrifugation at 5000 g was resuspended in 10 mM tris buffer (pH 8.0) and the protein solution was dialyzed against 10 mM tris buffer overnight at 4 ℃. α-Synuclein was purified by anion exchange chromatography (HiTrap Q HP; 20 mM tris buffer, pH 8) and size exclusion chromatography (HiLoad 16/60 Superdex 75 pg; 10 mM phosphate buffer, pH 7.4) at 4 ℃.

#### Tau

Recombinant full-length tau (2N4R) protein was expressed and purified from BL21(DE3) *E. coli* cells transformed with the pET15b plasmid (a gift from Dr. Smet-Nocca, Université de Lille, Sciences et Technologies, France) as previously described.^30^ Briefly, when the cells were grown at 37 °C in LB medium to an OD of 0.8, they were induced by addition of 0.5 mM IPTG and incubated for 3-4 hrs at 37 °C. After the induction, the cells were harvested by centrifugation. The bacterial pellet was resuspended in the lysis buffer and sonicated at 4 °C. The soluble fraction was heated at 80 ℃ for 20 min and the precipitates were removed by centrifugation. The supernatant containing tau protein was purified by cation exchange chromatography (HiTrap SP HP; 20 mM MES, 2 mM DTT, 1 mM MgCl_2_, 1 mM EGTA, 1 mM PMSF) followed by size exclusion chromatography (HiLoad 26/60 Superdex 200 pg; 10 mM phosphate buffer, 150 mM NaCl, 1 mM DTT, pH 7.4).

### Preparation of tau promoted α-synuclein filaments

To prepare α-synuclein filaments in the presence of tau, monomeric α-synuclein (70 μM in 10 mM phosphate buffer, pH 7.4) was mixed with tau monomers (20 μM in 10 mM phosphate buffer, pH 7.4) and incubated at 37 ℃ for 1 day under constant agitation at 250 rpm in an orbital shaker. Filamentous aggregates were examined with transmission electron microscopy (TEM).

### TEM

α-synuclein filamentous solution (1 mg/ml) was diluted by 20 times with 10 mM phosphate buffer (pH 7.4) and 5 μL of the diluted solution was placed on a formvar/carbon supported 400 mesh copper grid. After 30 sec incubation of the sample on the TEM grid, excess sample was blotted off with a filter paper. The grids were washed briefly with 10 μL of 1% uranyl acetate. The samples were then stained with 10 μL of 1% uranyl acetate for 30 sec and the excess stain was blotted off with a filter paper. The grids were then allowed to air dry and TEM images were collected using a Philips CM12 transmission electron microscope at an accelerating voltage of 80 kV.

### Cryo-EM data collection

A four microliter α-synuclein filamentous solution was applied to the back of each of the glow-discharged R2/1 Quantifoil grids. For the formation of vitrified ice, the grids were manually plunge-frozen into liquid nitrogen temperature cooled liquid ethane, after 3 seconds of blotting with filter papers. Grids were examined on Titan Krios electron microscope, equipped with GATAN K3 camera operated at 300kV. The defocus on camera was set to be randomly within 5,000-25,000Å range. The images have been collected with GATAN automated data collection software Latitude S (GATAN, Inc). The magnification was set to 81,000 and as a result the nominal pixel size is set to 1.1Å (the calibrated pixel size is found to be 1.07Å).

### Image processing

Movies were beam-induced motion corrected (in frame and among frames) and dose-weighted using MotionCor2^31^. Aligned (non-dose-weighted) integrated micrographs were used for contrast transfer function (CTF) estimation of each micrograph, using GCTF ^32^. Using Relion3-beta ^33^, filaments were manually picked and extracted with helical extraction. Two-dimensional (2D) classification in cisTEM ^34^ was performed to determine the segments with better quality and no crossing filaments. Segments from the best-looking classes were selected and moved to Relion for further processing. Helical 2D classification in Relion moved all of the segments into a limited number of classes, independent of the values used for regularization parameter (T). Using a cylinder (produced by relion_helix_toolbox) as the initial model and a very tight mask, the segments were helically 3D refined with local search for symmetry, starting from 4.7 Å and −1° values for helical rise and helical twist, respectively ^35^. The result of the refinement (resolution: ~ 8Å) was then lowpass filtered to 10 Å and then used for 3D classification without alignment with a very tight mask (T=25), into 6 classes which resulted in two improved classes with major portion of the particles (~243,000 and ~109,000 particles). Each of these two classes has been processed, but only the class with ~243,000 particles produced a higher resolution structure. Following the same methodology used in similar studies^36^, we continued with 3D classification into 1 class, starting with the class average from the last 3D classification lowpass filtered to 8 Å (T=35) with the same tight mask to focus the refinement on the separation of the subunits of the α-synuclein within the mask. Step by step increase of the value of T up to 45, resulted in a higher resolution for the reconstruction. Then to down-weigh the role of mask, we extended the binary mask much more beyond the diameter of the filament to include the structure inside the mask. Local search for helical twist and helical rise converged to −1.19° and 4.76Å. The structure hinted a higher-level helical symmetry with 179.36° and 2.43Å for helical twist and helical rise, respectively, and thus those values were used for further refinement. The handedness of the filaments was initially imposed arbitrarily. Later using a tomography data set of the same filament, the filaments were verified as left-handed. Auto-Refinement, with T=90, resulted in the best map with the highest resolution. Two separate rounds of beam-tilt correction using CtfRefine in Relion were done to improve the overall resolution as well as the map visual quality. Per particle CTF refinement, however, did not improve the resolution of the map. Using Relion post-processing, we were able to determine the overall reconstruction resolution to be 4 Å (Figure S1). Local resolution was determined using the corresponding Relion tool and local sharpening was done using LocalDeblur^37^. Over-sharpening was seen in last iterations of local sharpening. Consequently, the result with the lowest amount of noise was selected for model building.

### Atomic model building and refinement

Resolution of the tau-promoted α-synuclein filament density map was not high enough for de novo modeling of the structure. However, our solid-state NMR data showed that the resonances for certain residues in our filaments are similar to those of a previously reported structure (PDB: 6rt0) for α-synuclein filaments. Hence, the atomic model was built, using 6rt0 as the initial model, starting from the region having residues with resonances of high similarity (residues 53-65) in Coot^38^. Then a poly-alanine model was built into the density and the residues were later replaced by the correct sequence. The atomic model was refined using Phenix real space refinement ^39^, manually modified in Coot ^38^ and validated using Phenix ^40^ (Table S1 and S2).

Lack of well-resolved sidechains makes it difficult to investigate salt-bridges between protofilaments. Therefore, we used MDFF ^41^ with explicit solvent to look at the molecular dynamic interactions in the atomic model. Water molecules were added in VMD ^42^ and the solvent was neutralized with 150 mM of NaCl to simulate physiological ionic strength conditions. Salt-bridges between protofilaments (K45, E46) were detected using the Salt-bridge module of VMD.

#### Data availability

The electron density map is available in Electron Microscopy Data Bank (EMDB) with ID EMD-23212 and the atomic model is available in Protein Data Bank (PDB) with ID 7L7H. The raw data, intermediate maps, masks, and intermediate atomic models are all available from the authors upon request.

## Results

### Solid-state NMR of Tau-promoted α-synuclein filaments

Monomeric α-synuclein (70 μM) was incubated in the presence of tau monomers (20 μM) at 37 ℃ in 10 mM phosphate buffer (pH 7.4). Long homogeneous filamentous aggregates were observed after 24 hrs of incubation in the presence of tau (Figure 2). Solid-state NMR was initially used to compare structural features of the tau-promoted filaments to those of previously reported in vitro α-synuclein filaments (Figure 3). The two-dimensional ^13^C-^13^C correlation spectrum obtained with dipolar-assisted rotational resonance mixing scheme (DARR)^43^ suggests that the tau-promoted filaments (Figure 3a) have distinct molecular conformations from those of two α-synuclein filaments (Figure 3c and 3d). On the contrary, the 2D DARR spectrum of the tau-promoted filaments is somewhat similar to that of the in vitro filament (red in Figure 3b) with notable differences (black in Figure 3b). These solid-state NMR results suggest that the co-factor, tau, appears to induce the formation of a specific fibrillar conformation.

**Figure 2.**
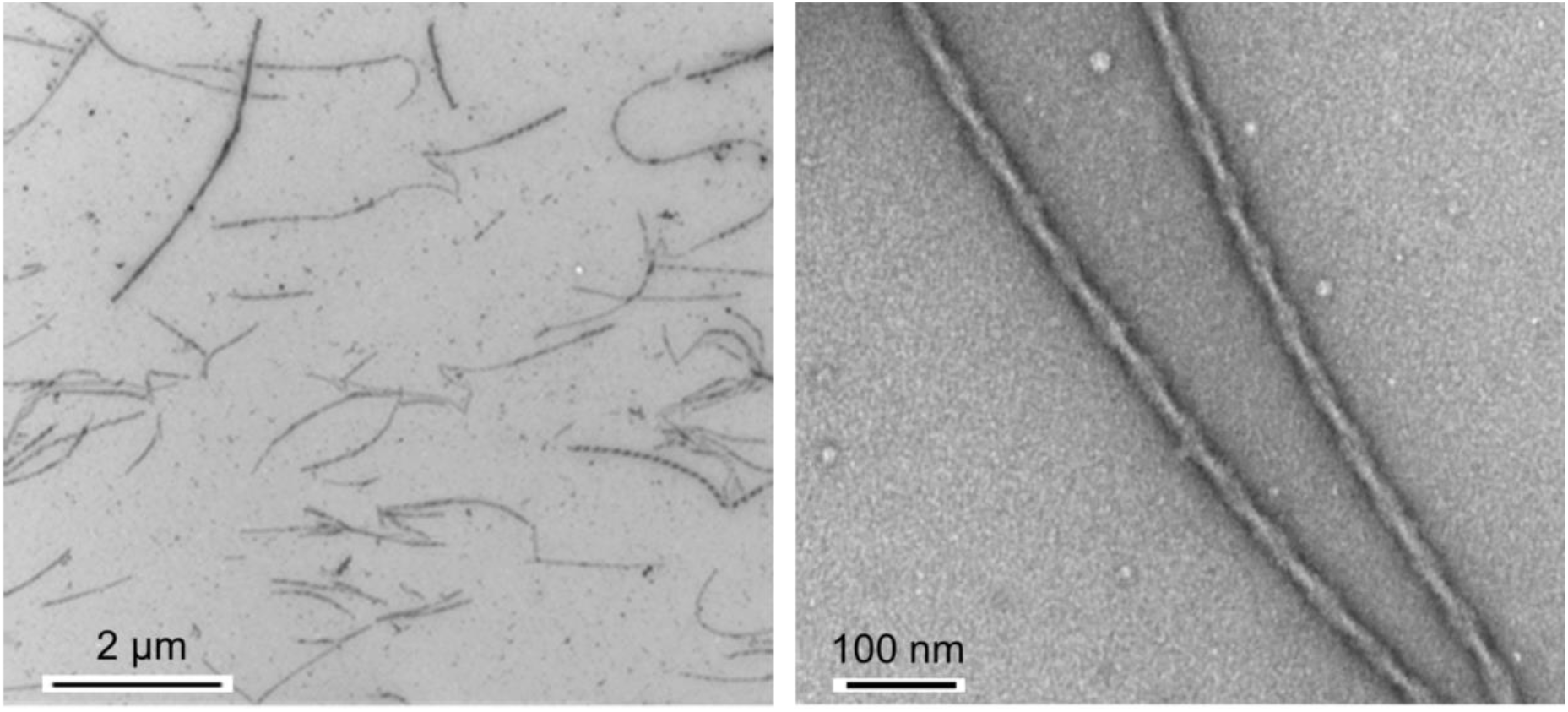
Representative TEM images of tau-promoted α-synuclein filaments showing the homogeneous twisting filaments.

**Figure 3.**
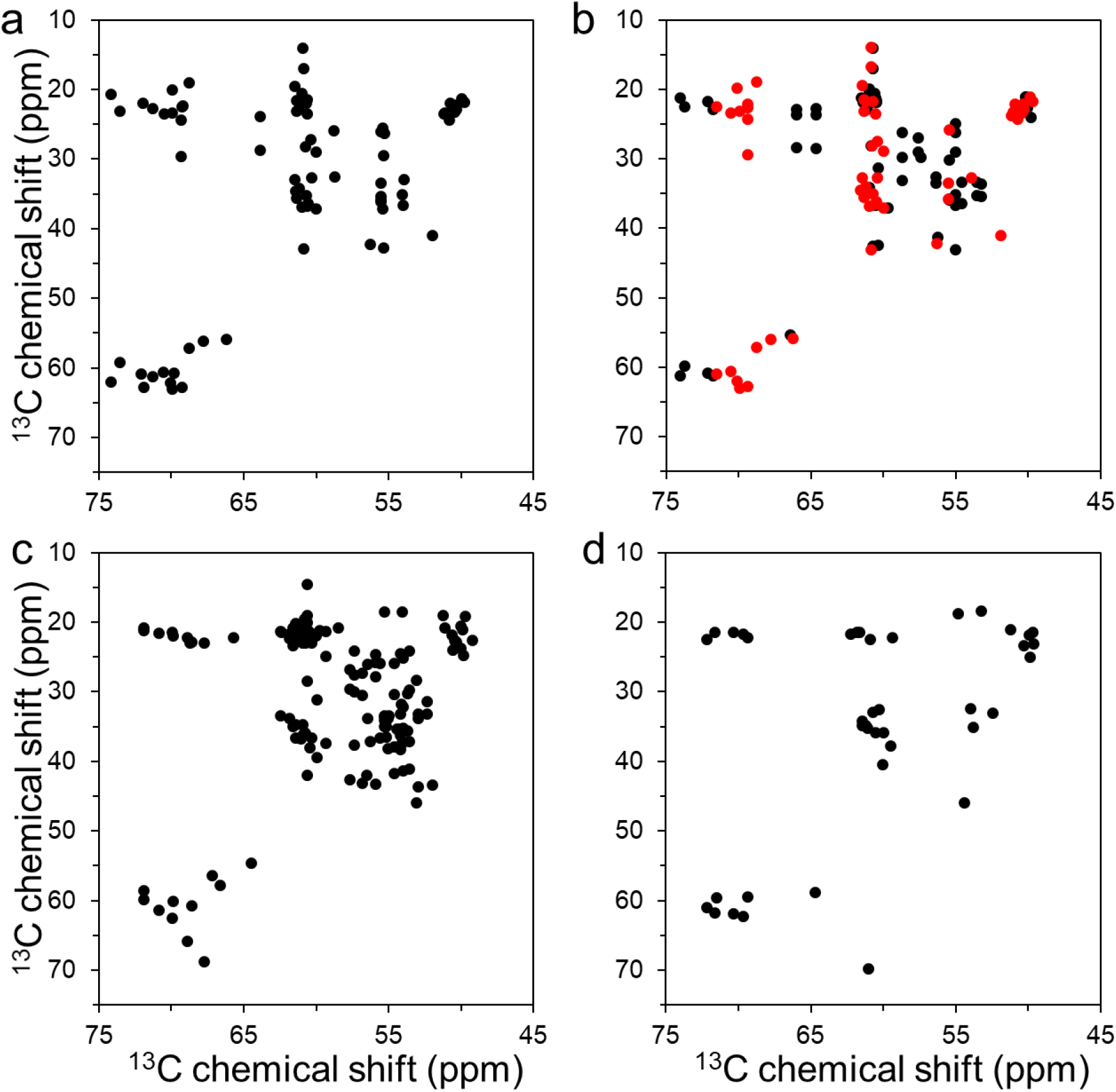
Overview of aliphatic region of 2D ^13^C-^13^C DARR NMR spectra of uniformly ^13^C/^15^N labeled α-synuclein filament polymorphs. (a) Tau-promoted α-synuclein polymorph. (b) Fibril-type α-synuclein polymorph (BMRB 18860)^44^. (c) Ribbon-type α-synuclein polymorph (BMRB 17498).^45^ (d) Greek-key type α-synuclein polymorph (BMRB 25518)^16^. Cross-peaks with similar NMR resonances for the tau-promoted α-synuclein polymorph and ribbon-type polymorph are colored red in 3b. The ribbon- and fibril-type polymorphs of α-synuclein have distinct molecular packing arrangement and intermolecular interactions.^44, 46^ The NMR cross-peaks were drawn using our experimental DARR spectrum for the tau-promoted filaments (a) and chemical shifts reported in BMRB for the previously reported DARR spectra of α-synuclein filaments (b – d).

### Cryo-EM structure of the tau-promoted α-synuclein filaments

Cryo-EM was then used to determine near-atomic structure of the tau-promoted α-synuclein filaments. Preformed tau-promoted filaments were frozen on a carbon-coated grid and images were acquired at 81,000X magnification on a Titan Krios (300 kV) equipped with a K3 GATAN direct electron detector camera. About 240,000 segments extracted from 1,800 micrographs were analyzed using Relion reference-free two-dimensional (2D) classification. The initial classification analyses revealed one major species in the 2D classes (Figure 4a). The 2D classes show that the protofilaments are twisted around with a crossover distance of 610 Å (Figure 4b) and a helical rise of 4.8 Å based on the power spectrum (Figure 4c). The left-twisting handedness was determined by cryo-electron tomography. The 2D classes were used for three dimensional (3D) helical reconstruction in Relion 3, which resulted in a 3D density map at 4.0 Å resolution (Figure 5a).

**Figure 4.**
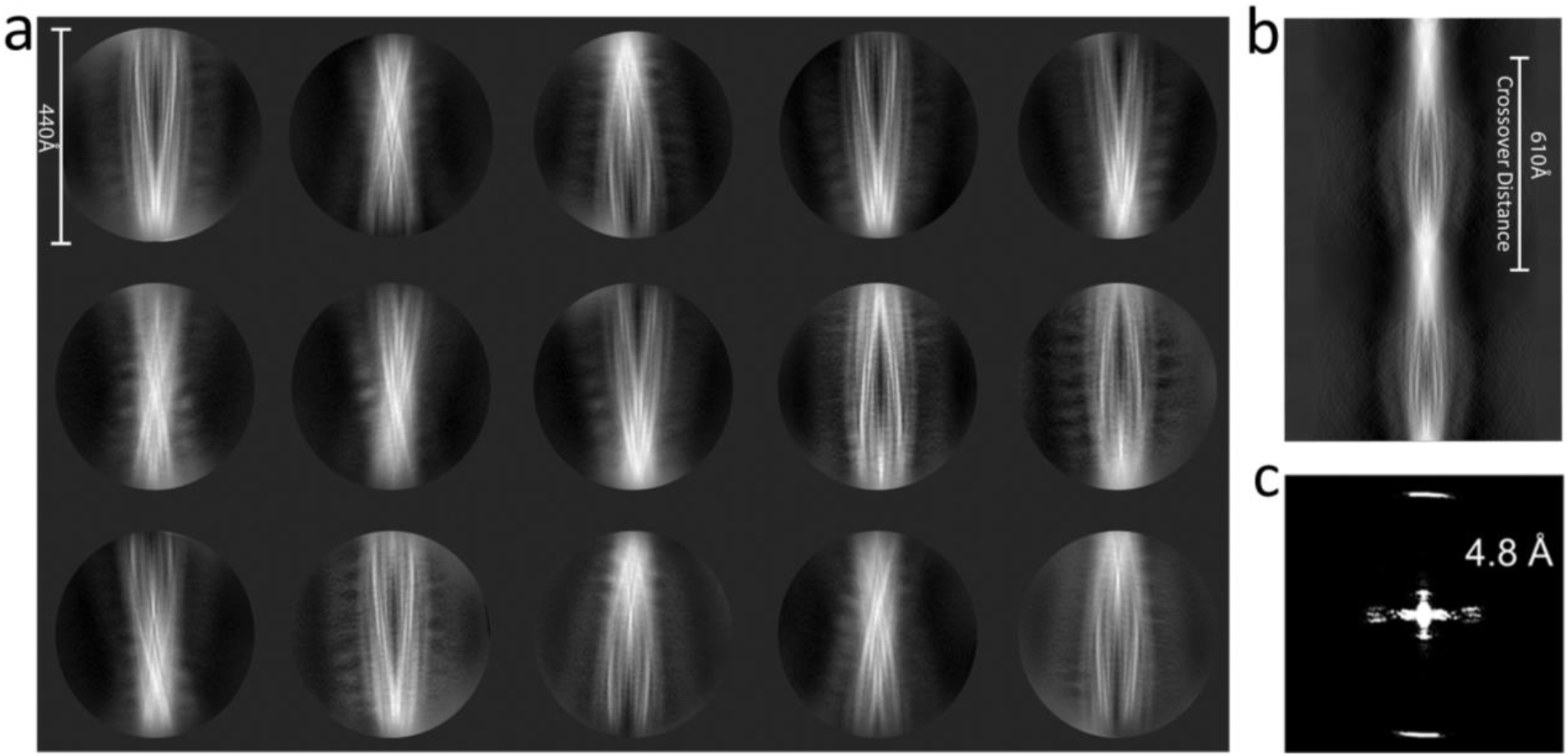
2D class averages of the tau-promoted α-synuclein filaments. (a) Representative 2D class averages of tau derived filaments using cisTEM (box size of 440 Å). (b) A sinogram representing the full rotation along the helical axis of the filament produced by relion_helix_inimodel2d as described by Scheres.^47^ (c) The power spectrum of selected 2D reference-free class averages.

**Figure 5.**
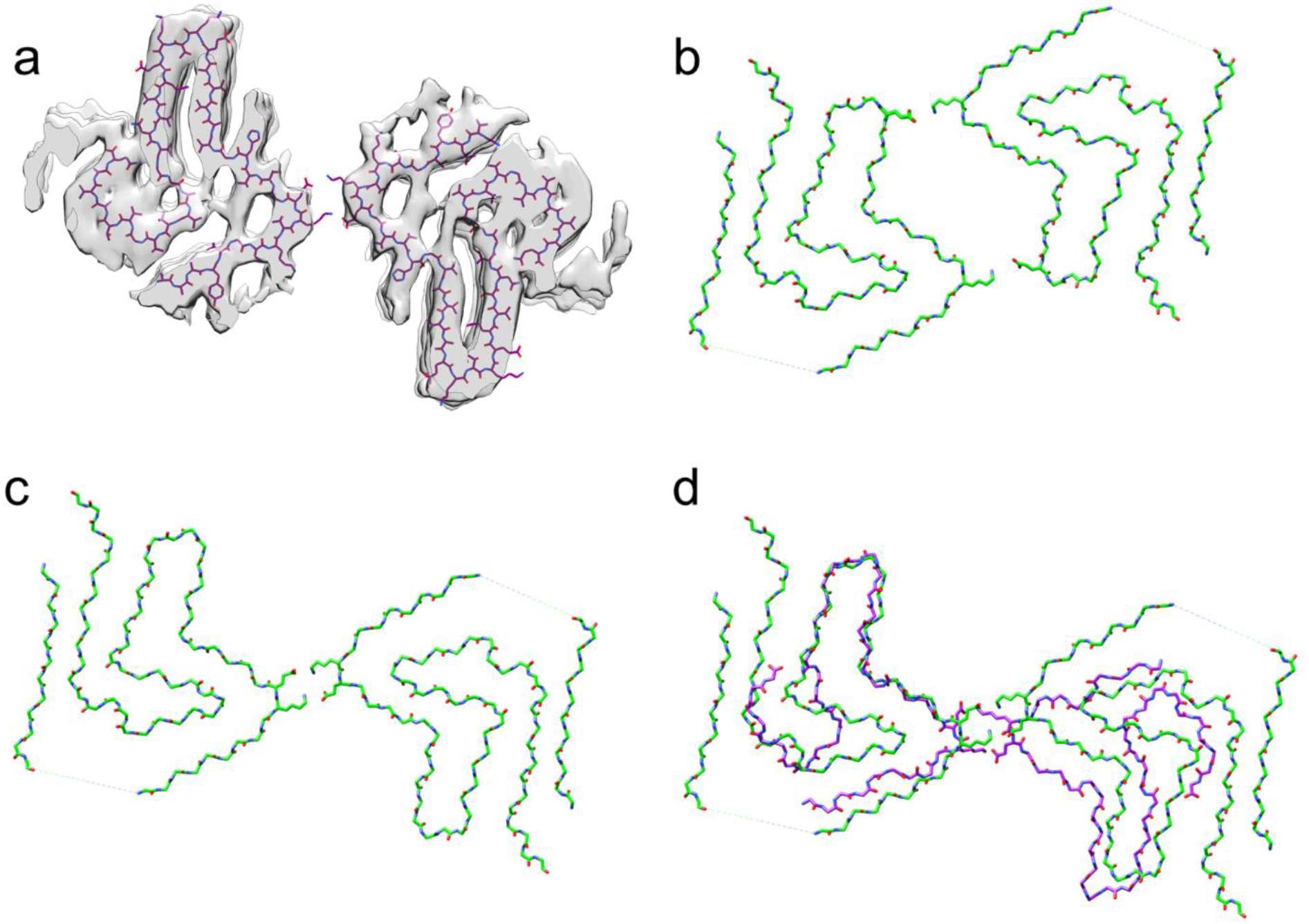
Structural comparison of α-synuclein filament polymorphs. (a) Overlay of the tau-promoted α-synuclein filament atomic model on the density map. (b and c) α-synuclein filament polymorphs 2a and 2b, respectively, determined by previous cryo-EM structural studies.^20^ (d) Overlaid structures of the tau-promoted α-synuclein filament (purple) and polymorph 2b (green). The same salt bridge between the residues K45 and E46 was observed in the interfacial region of the polymorph 2b (Figure 5c) and tau-promoted α-synuclein polymorph (Figure 5d).

The 3D density map of the tau-promoted α-synuclein filaments was similar to that of the polymorph 2b, as was suggested by our solid-state NMR (Figure 3a and 3b). It is, however, interesting to note that tau induced the formation of only one polymorph (Figure 5c), although the previous study showed that the protofilaments were assembled into the two fibril polymorphs in the same buffer (Figure 5b and 5c).^20^ The tau-promoted α-synuclein filament also exhibits notable differences in comparison to that of the polymorph, particularly the N- and C-terminal regions (Figure 5d and Figure S2). Firstly, the interaction between the N-terminal (15-20) and C-terminal (85-91) regions are not observed in the tau-promoted filament. Secondly, the more extensive C-terminal region (80-140) is disordered in comparisons with that of the other structure (91-140), which might be due to interactions between the positively charged tau and negatively charged C-terminal region of α-synuclein (Figure S2). Thirdly, the tau-promoted filaments with a half-pitch of 63 nm are twisted much faster in comparison with that of polymorph 2b (96 nm) (Figure 5d and Figure S3).

The structural model for the tau-promoted filaments was compared to the previously reported structures of α-synuclein filaments (Figure 6). Our tau-promoted α-synuclein filaments adopt an overall Greek-key type structure observed in the first solid-state NMR structure of α-synuclein filaments.^16^ However, several regions including the N- and C-terminal regions (residues 36-48 and 66-79) are notably different from the previous Greek-key type structures, as was suggested by our solid-state NMR results (Figure 3a and 3d). In addition, interfacial contacts between the two protofilaments and the degree of helical twist (Table S3) are quite distinct from those of the previously reported structures. These results indicate that interactions between co-factors and α-synuclein may lead to distinct molecular conformations and intermolecular contacts between the protofilaments of α-synuclein.

**Figure 6.**
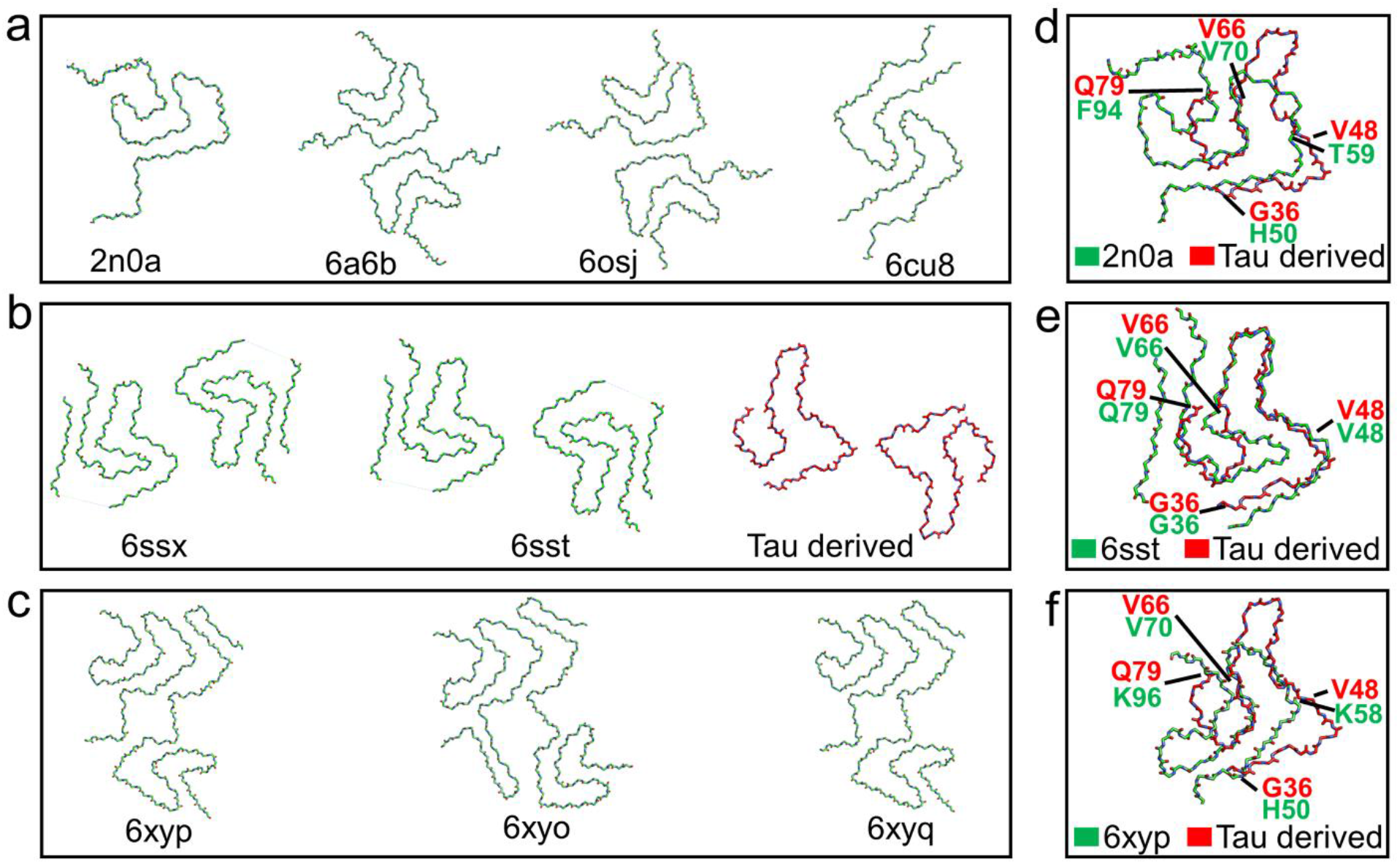
Structural comparison of various polymorphs of full-length α-synuclein filaments. Representative structures of (a) α-synuclein polymorphs 1a (PDB 2n0a, 6a6b)^16, 17^ and polymorph 1b (PDB 6cu8)^18^. (b) α-synuclein polymorph 2a (PDB 6ssx)^20^, polymorph 2b (PDB 6sst)^20^ and tau-promoted α-synuclein polymorph (this study, PDB 7l7h). (c) MSA patient derived α-synuclein polymorph type-1 (PDB 6xyp)^23^ and type-2 (PDB 6xyo, 6xyq)^23^. (d-f) Overlay of protofilament folds of tau-promoted α-synuclein filament with polymorphs 1, 2 and ex vivo MSA polymorphs, respectively.

## Discussion

Molecular mechanism by which α-synuclein self-assembles into fibrillar aggregates in vivo has remained largely unknown. It was previously shown that monomeric α-synuclein is stabilized by long-range interactions between the N- and C-terminal regions.^48, 49^ Perturbations of the long-range interactions may initiate misfolding and aggregation of α-synuclein. Indeed, various co-factors that interact with the N- and/or C-terminal regions promoted the formation of fibrillar aggregates of α-synuclein.^50–52^ Recently solved cryo-EM structures of ex vivo α-synuclein filaments extracted from MSA and DLB patients revealed that the ex vivo filaments adopt distinct molecular structures from those of in vitro α-synuclein filaments,^23^ supporting that co-factors may play important roles in promoting α-synuclein aggregation in vivo. Comparative structural analyses of α-synuclein aggregates derived by co-factors and ex vivo aggregates will, therefore, be required to identify co-factors that may play critical roles in α-synuclein aggregation in vivo.

Several lines of evidence indicate that pathological proteins such as β-amyloid (Aβ) peptides, tau and α-synuclein synergistically promote their mutual aggregation.^24, 53–61^ In particular, co-existence of tau and α-synuclein aggregates in synucleinopathy patient’s brains suggests that tau may interact with α-synuclein, accelerating the formation of fibrillar α-synuclein aggregates in vivo. In this work, we solved cryo-EM structure of α-synuclein filaments derived by tau and compared the structure to those of previously reported structures of α-synuclein filaments. Previous structural studies of α-synuclein filaments revealed that α-synuclein can form diverse filamentous aggregates with distinct molecular conformations (Figure 6). Polymorphic structures were also observed for the filaments formed even in the same buffer.^16, 18, 20^ It is plausible that multiple conformers in the conformational ensemble of disordered α-synuclein are able to form diverse α-synuclein filaments with different molecular conformations and/or different interfaces between the protofilaments (Figure 1). Interestingly, tau-promoted α-synuclein filaments adopt a Greek-key type structure similar to one of the polymorphic α-synuclein filaments. However, the detailed molecular conformation and the degree of the helical twist are different from those of the polymorphs (Table S3). In addition, recent studies revealed that poly(ADP-ribose) may interact with α-synuclein in vivo^62^ and induce the formation of a more toxic α-synuclein strain with distinct molecular conformations.^25^ These results suggest that the interaction between co-factors and α-synuclein may direct the protein to a specific misfolding and aggregation pathway toward the distinct α-synuclein filament, highlighting the importance of cellular environments in protein misfolding and aggregation.^63^

Previous structural studies of the full-length and truncated α-synuclein filaments revealed that the C-terminally truncated α-synuclein filaments have increased helical twists even though the full-length and truncated filaments adopt almost identical core structures,^19, 64^ suggesting that the negatively charged C-terminal region affects the helical twist in the parallel alignment. Thus, the increased helical twist of the tau-promoted full-length α-synuclein filaments may result from the electrostatic interaction between the positively charged tau and negatively charged C-terminal region of α-synuclein, which may reduce repulsive interactions between the C-terminal regions in the parallel alignment and facilitate the tighter helical twist. The longer disordered C-terminal region (residues 81 – 140) in the tau-promoted filament compared to that of the previously reported α-synuclein filaments (residues 90 – 140) may also result from the interaction between the tau and C-terminal regions of α-synuclein.

In summary, we report a distinct molecular structure of the α-synuclein filament formed in the presence of tau. The interaction between the C-terminal region of α-synuclein and tau leads to a distinct molecular conformation of α-synuclein filament with a shorter helical pitch. These results suggest that interaction between α-synuclein and various potential co-factors in cellular environments may promote the formation of diverse α-synuclein filaments with different molecular conformations. More extensive comparative structural analyses of in vitro α-synuclein filaments derived by co-factors and ex vivo α-synuclein filaments extracted from the patients are required to better understand molecular mechanism of α-synuclein aggregation in vivo.

## ASSOCIATED CONTENT

### Supporting Information

FSC curve. Structural comparison of tau derived α-synuclein polymorph and polymorph 2b. Density maps showing the helical twisting patterns α-synuclein polymorph and polymorph 2b. Cryo-EM data collection, refinement, and validation statistics. Helical twists comparison of various α-synuclein polymorphs.

The following files are available free of charge.

## AUTHOR INFORMATION

### Author Contributions

The manuscript was written through contributions of all authors. All authors have given approval to the final version of the manuscript.

### Notes

The authors declare no competing financial interest.

## ACKNOWLEDGMENT

We thank Dr. Jun-yong Choe (East Carolina University) for helpful discussion. We also thank Hamidreza Rahmani for helpful suggestions on molecular dynamics analysis on the atomic model.

## ABBREVIATIONS

NMR: nuclear magnetic resonance
TEM: transmission electron microscopy
DARR: dipolar assisted rotational resonance
cryo-EM: cryo-electron microscopy

## Accession Codes

α-synuclein: UniProtKB entry P37840

tau: UniProtKB entry P10636

## For Table of Contents use only

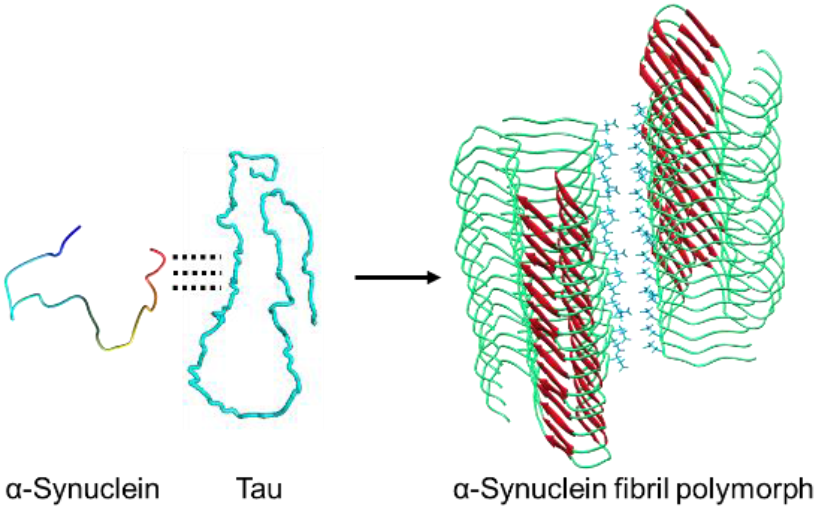

