## Supporting Information for "Distinct cryo-EM Structure of α-synuclein Filaments derived by Tau"

[1]Institute of Molecular Biophysics, Florida State University, Tallahassee, FL 32306-4380, USA. [2]Department of Chemistry, East Carolina University, Greenville, NC 27858, USA. [3]Departments of Neurology, Neuroscience and Cell Biology, University of Texas Medical Branch, Galveston, TX, 77555, USA. [4]Genome Integrity and Structural Biology Laboratory, National Institute of Environmental Health Sciences, National Institutes of Health, Department of Health and Human Services, Research Triangle Park, NC, 27709, USA. [5]Department of Chemistry and Francis Bitter Magnet Laboratory, Massachusetts Institute of Technology, Cambridge, MA, 02139, USA.

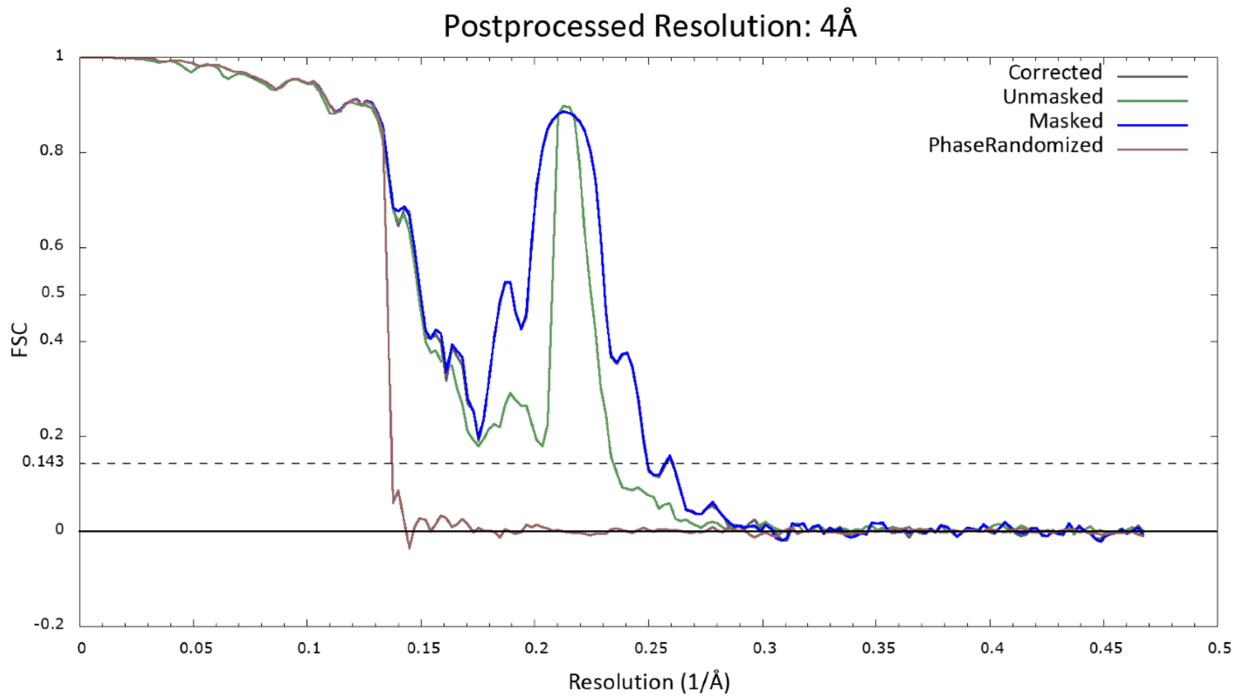

**Figure S1.** Gold-standard Fourier Shell Correlation (FSC, 0.143 criterion) curve of the two half-maps, produced in Relion 3.1, demonstrating an estimate of the global resolution of the map to be 4 Å.

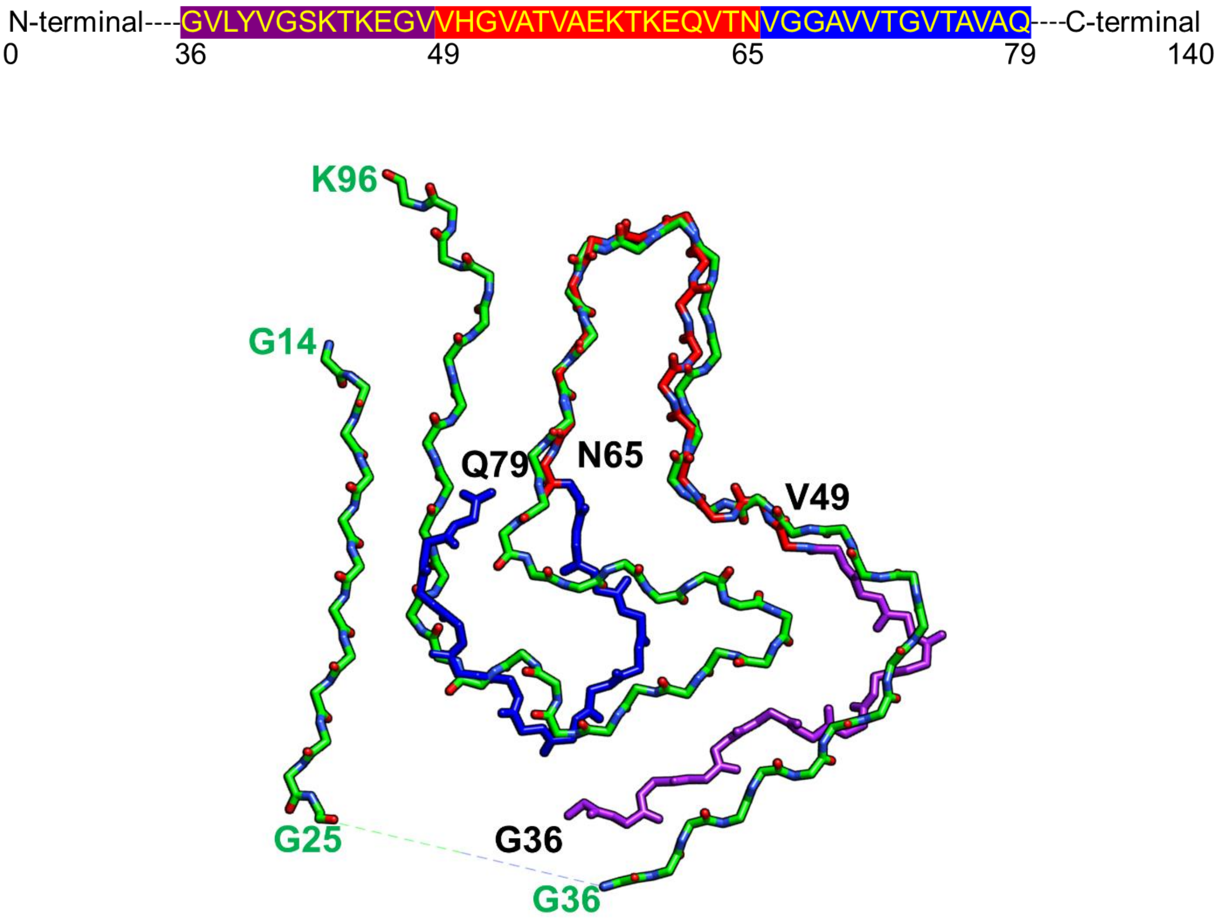

**Figure S2.** Overlaid structures of tau-promoted α-synuclein polymorph (purple) and polymorph 2 (green) highlighting the structural differences between the two polymorphs. Residues (49-65) with similar NMR resonances for tau derived polymorph and polymorph 2 are colored in red.

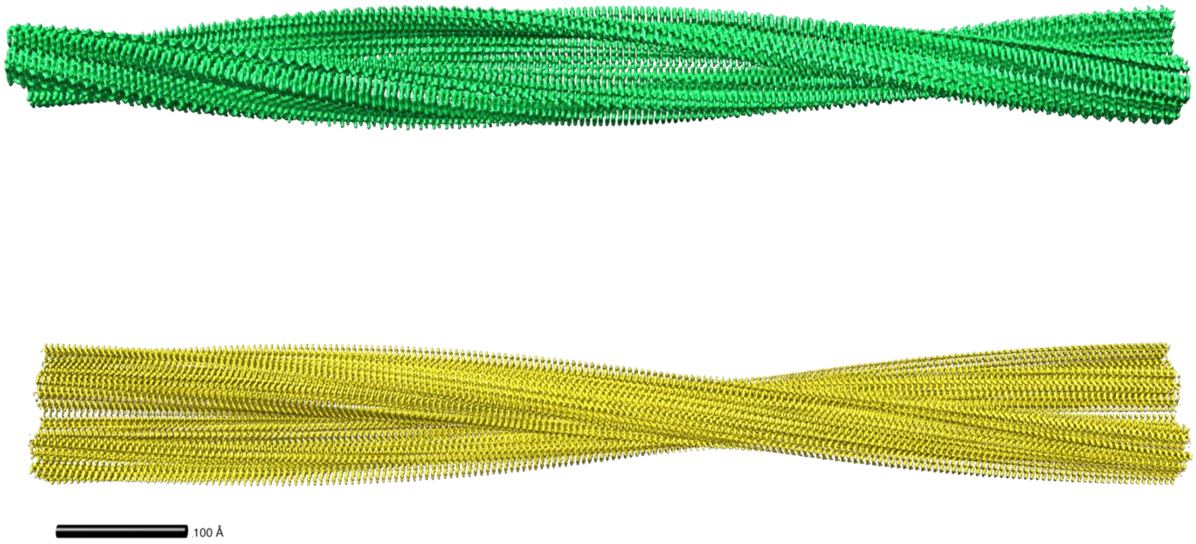

**Figure S3.** Comparison of degree of helical twist for tau-promoted α-synuclein polymorph (top, green) and polymorph 2b (bottom, yellow) showing the tau-promoted α-synuclein polymorph twists faster.

**Table S1.** Cryo-EM data collection and reconstruction statistics of the tau-promoted α-synuclein fibril.

| Cryo-EM statistics of tau derived α-synuclein fibril (PDB: 7l7h) | |
|---|---|
| **Data Collection** | |
| Camera | K3 |
| Magnification | 81K |
| Physical pixel size | 1.11Å |
| Calibrated pixel size | 1.074Å |
| Defocus range (μm) | -0.5 to -2.5 |
| Voltage (kV) | 300 |
| Number of Frames | 45 |
| Dose per frame (e/A$^2$) | 1 |
| **Reconstruction** | |
| Box size (pixels) | 400 |
| Inter-box distance (Å) | 20 |
| Number of Micrographs | 1,799 |
| Number of filaments | 33,726 |
| Initial number of segments | 1,043,263 |
| segments after 2D classification (cisTEM) | 352,578 |
| segments after Relion 3D classification | 243,181 |
| Final number of segments | 243,181 |
| Map overall resolution (Å) (refinement) | 4.0 |
| FSC threshold | 0.143 |
| Map resolution range (Å) (MonoRes) | 3.75 to ~50.0 |
| Helical rise (Å) | 2.43 |
| Helical twist (degree) | 179.3 |

**Table S2.** Cryo-EM model building statistics of the tau-promoted α-synuclein fibril.

| Cryo-EM validation report of tau derived α-synuclein fibril | |
|---|---|
| **Atomic model** | |
| Initial model (PDB code) | 6rt0 |
| Model Composition | |
| Atoms (Hydrogens) | 2439 (0) |
| Protein residues | 352 |
| Ligands | 0 |
| R.M.S deviation | |
| Bond lengths (Å) (# > 4σ) | 0.008 (0) |
| Bond angles (°) (# > 4σ) | 1.473 (8) |
| Validation | |
| MolProbity score | 2.94 |
| Clashscore | 13.43 |
| Poor rotamers (%) | 3.53 |
| Ramachandran Plot | |
| Favored (%) | 63.69 |
| Allowed (%) | 36.31 |
| Disallowed (%) | 0.0 |

**Table S3.** Comparison of degree of helical twists (half-pitch) for α-synuclein polymorph 1a (6a6b, 6osj, 6cu7), polymorph 2a (6ssx), polymorph 2b (6sst) and tau-promoted polymorph (7l7h).

| PDB ID | Half-pitch (nm) |
|---|---|
| 6a6b | 120 |
| 6osj | 121 |
| 6cu7 | 92 |
| 6ssx | 108 |
| 6sst | 96 |
| Tau derived (7l7h) | 63 |
